# Assessing fish biodiversity and community structure in the Ganjiang River using eDNA metabarcoding

**DOI:** 10.64898/2026.10.08.753657

**Authors:** Zeguo Zeng, Chanjuan Qu, Jinwei Lv, Ming Zhang, Jianjun Li, Qin Guo, Hai Lian, Jing Guo, Yuxiang Wang, Qingxiang Zeng, Binsong Jin

## Abstract

Hydropower development has emerged as a major driver of global environmental change, causing significant ecological damage to freshwater ecosystems and threatening aquatic biodiversity. However, the impact of cascade dams on fish communities in large river systems remains poorly understood, particularly in biodiversity hotspots such as China. Here, we used environmental DNA (eDNA) metabarcoding to assess the spatiotemporal patterns of fish diversity and community structure in the Ganjiang River, the largest tributary of the Yangtze River, in relation to cascade hydropower development. We collected water samples from 19 sites along the main stem and major tributaries in both wet and dry seasons, and analyzed the eDNA data using high-throughput se-quencing and advanced bioinformatics pipelines. Our results revealed significant seasonal and spatial variations in fish diversity and community composition, with lower species richness and distinct assemblages in the impounded sections upstream of the dams compared to the downstream reaches. The fish communities were strongly influenced by environmental factors such as water temperature, pH, dissolved oxygen, and nutrients, which exhibited complex interactions with dam operation and land use practices. Spatial scale analyses further indicated that the impact of dams on fish diversity was most pronounced at the local scale, but also detectable at the regional scale. Our findings provide novel insights into the ecological consequences of cascade hydropower develop-ment on fish communities and highlight the importance of considering spatial scales and envi-ronmental heterogeneity in assessing the impacts of human activities on freshwater biodiversity. Our study demonstrates the power of eDNA metabarcoding as a cost-effective and non-invasive tool for biodiversity assessment and conservation planning in large river systems facing multiple stressors and knowledge gaps.

## 1. Introduction

Hydropower development has emerged as a significant driver of global environmental change, with China being a major hotspot of dam construction (Sun et al., 2019). While these projects provide important socio-economic benefits, the rapid proliferation of large dams has led to substantial ecological damage to freshwater ecosystems, including the the decline and extinction of numerous endemic fish species in major rivers such as the Yangtze River (Cheng et al., 2015; Xu et al., 2023). As hydropower development continues to expand, there is an urgent need to assess and mitigate its impact on fish diversity in China’s rivers.

The Ganjiang River, the largest tributary of the Yangtze River, is a critical watershed for freshwater fish conservation in southeastern China (Guo et al., 2018). Historical records have documented 118 fish species in this river system, including many endemic and threatened ones (Guo et al., 2018; Hu et al., 2019). However, like many other rivers in China, the Ganjiang River has been heavily fragmented by a cascade of six large dams since the 1990s (Hu et al., 2019). The impact of these dams on fish diversity and community structure remains poorly understood, highlighting the need for comprehensive assessments to inform conservation and management strategies.

Fish are essential components of riverine ecosystems, playing vital roles in maintaining biodiversity, food web dynamics, nutrient cycling, and ecosystem functioning (Gao et al., 2023). Despite covering only 0.8% of the Earth’s surface, freshwater ecosystems harbor more than 40% of all fish species (Dudgeon et al., 2006). However, this extraordinary diversity is increasingly threatened by anthropogenic stressors, including habitat degradation, pollution, overfishing, invasive species, and climate change (Reid et al., 2019). Among these threats, dam construction stands out as one of the most pervasive and damaging to freshwater fish worldwide (Chen et al., 2023). Dams fragment river systems, disrupt hydrological connectivity, alter natural flow regimes, and create barriers for fish movements and migrations, leading to cascading effects on fish populations and communities (Limburg and Waldman, 2009; Spinti et al., 2023; Timpe and Kaplan, 2017). These impacts are often exacerbated when multiple dams are built in a cascade manner along the same river (Winemiller et al., 2016).

Traditionally, fish diversity is assessed by capture-based methods such as gillnetting, electrofishing, and angling. While providing valuable information, these methods have limitations in terms of labor intensity, cost, invasiveness, and detection bias (Bonar et al., 2017). Environmental DNA (eDNA) metabarcoding has emerged as a promising tool to overcome these challenges, enabling rapid, non-invasive, and high-resolution assessment of fish diversity in various aquatic ecosystems (Norgaard et al., 2021; Sahu et al., 2023). By analyzing the genetic material shed by organisms into water, eDNA metabarcoding allows for efficient and sensitive detection of fish species across multiple taxonomic groups and habitat types (Ruppert et al., 2019; Stat et al., 2017).

In this study, we employed eDNA metabarcoding to investigate the spatiotemporal patterns of fish diversity and community structure in the Ganjiang River, focusing on the impact of cascade hydropower development. We collected water samples from 19 sites along the main channel and major tributaries in both wet and dry seasons, and analyzed the eDNA data using high-throughput sequencing and bioinformatics. Our main objectives were to: (1) assess the spatial-temporal patterns of fish diversity based on eDNA; (2) compare the fish community structure and ecological groups among different river sections in relation to dam locations; and (3) evaluate the impact of dams on fish biodiversity and provide conservation implications. To our knowledge, this is the first comprehensive assessment of fish diversity in the Ganjiang River using eDNA metabarcoding, and one of the few studies to evaluate the ecological consequences of cascade damming on fish communities in large subtropical rivers. By integrating molecular, taxonomic, and ecological approaches, we aim to provide novel insights into the biodiversity impacts of hydropower development in the Yangtze Basin, and inform future conservation, monitoring, and management strategies for the rich fish fauna in the Ganjiang River and beyond. Our findings also highlight the potential of eDNA metabarcoding as a powerful tool for aquatic biodiversity assessment in the era of big data and global change.

## 2. Materials and methods

### 2.1. Study area and sample collection

#### 2.1.1. Selection of sampling sites

To assess the impact of dam construction on fish diversity in the Ganjiang River, we selected 19 sampling sites along the main channel, covering both the free-flowing sections and the impounded sections (Fig. 1). The sites were chosen based on their representativeness of different habitat types (e.g., riffles, pools, runs, and backwaters), spatial distribution in relation to the six large dams (Wan’an, Jingangshan, Shihutang, Xiajiang, Xingan, and Longtoushan) and major tributaries (e.g., Yuanshui River, Lushui River, Minjing River), and accessibility and safety conditions for field sampling (Cilleros et al., 2019). This site selection strategy aimed to capture the spatial heterogeneity of fish communities and environmental conditions in the river network, as well as the potential influence of dam-induced habitat fragmentation and flow regulation (Agra et al., 2023). Among the 19 sites, 11 were located in the free-flowing sections above or between the dams, and 8 were located in the impounded sections within the reservoirs. The distance between adjacent sites ranged from 10 to 50 km, with an average of 30 km. The elevation of the sites ranged from 20 to 200 m above sea level.

**Figure 1.**
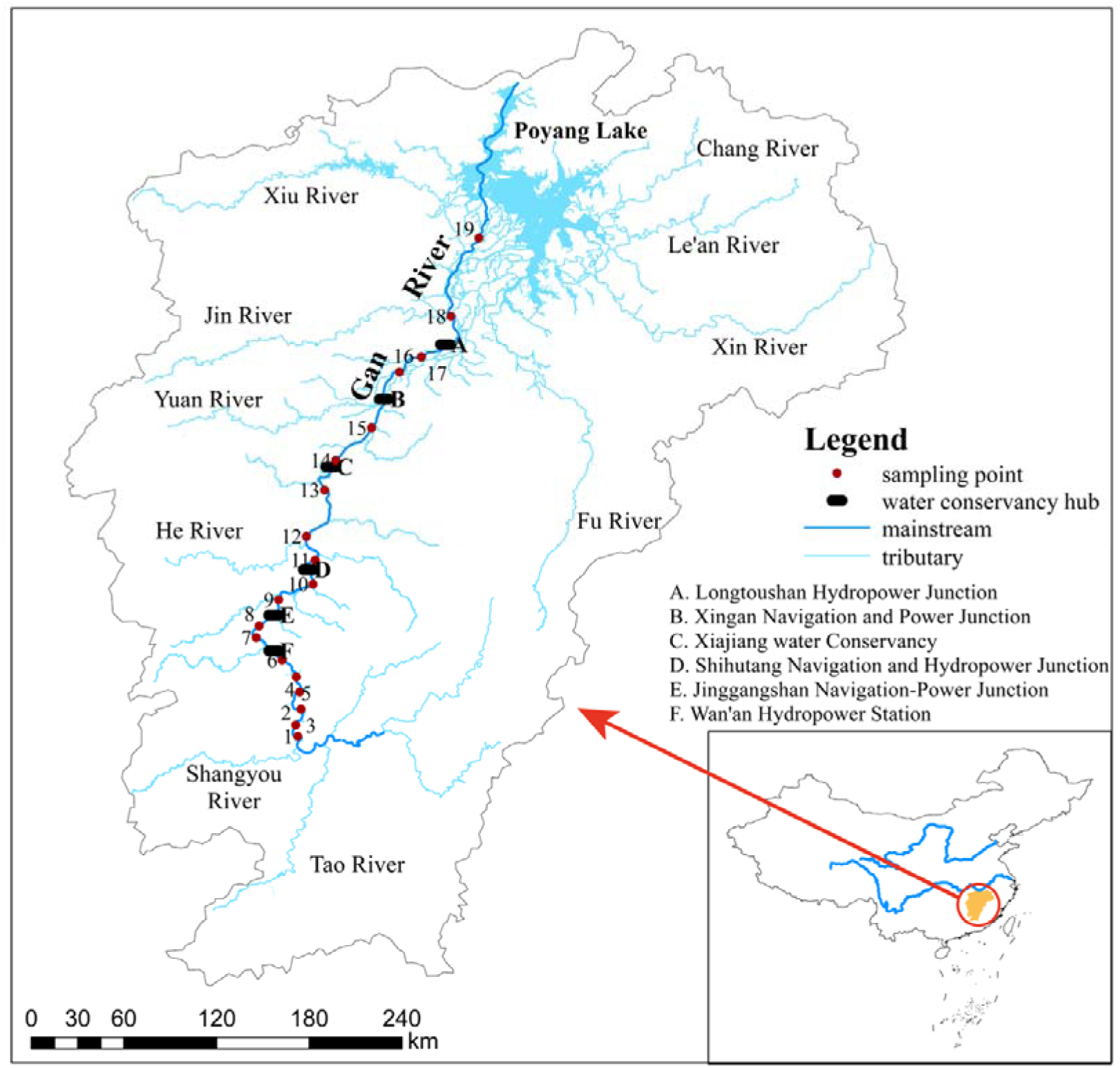
Map of the Ganjiang River basin showing the locations of sampling sites and cascade dams.

#### 2.1.2. Water sample collection and processing

We collected water samples for eDNA analysis at each site in both wet (July 2021) and dry (December 2021) seasons, with all 19 sites sampled within one week in each season to minimize temporal variation. At each site, three replicate 1-L water samples were collected from the subsurface layer at the left bank, right bank, and middle of the channel using a sterile polypropylene bottle attached to a telescopic pole. The bottle was rinsed three times with river water before sample collection. In addition, we measured environmental variables, including water temperature, pH, dissolved oxygen, conductivity, turbidity, total nitrogen, total phosphorus, and chlorophyll a, using a multiparameter water quality sonde (YSI EXO2, USA) to characterize habitat conditions and analyze factors influencing fish community structure.

The water samples were immediately filtered through 0.45-μm cellulose nitrate membranes using a portable vacuum pump. The filtration equipment was sterilized with 10% bleach solution and rinsed with deionized water between samples to prevent cross-contamination. Each filter was placed in a tube with 95% ethanol and stored in a cooler until being transported to the laboratory, where they were stored at −20°C until DNA extraction. Negative control samples were processed in parallel to monitor potential contamination during sampling and lab work (Sales et al., 2019).

### 2.2. eDNA extraction, PCR amplification, and sequencing

#### 2.2.1. DNA extraction and purification

The eDNA was extracted from the filter membranes using the DNeasy PowerWater Kit (Qiagen, Germany) following the manufacturer’s protocol. The extracted DNA was assessed for purity and quantity using a NanoDrop spectrophotometer and a Qubit fluorometer, respectively. Samples with low quality or yield were re-extracted or concentrated before further processing.

#### 2.2.2. PCR amplification and sequencing library preparation

A two-step PCR approach was used to amplify and index the fish eDNA. In the first step, the mitochondrial 12S rRNA gene was amplified using the MiFish universal primer set(Miya et al., 2015; Xiong et al., 2022). The PCR products were purified and then indexed with unique dual barcodes using the Nextera XT Index Kit in the second step. The indexed libraries were quantified, normalized, pooled, and sequenced on an Illumina MiSeq platform to generate 2 × 300 bp paired-end reads.

#### 2.2.3. Sequencing data processing and taxonomic assignment

The raw sequencing data were processed using the DADA2 pipeline in QIIME 2 to filter, denoise, and dereplicate the reads into amplicon sequence variants (ASVs). The ASVs were then classified taxonomically using the BLAST+ consensus classifier against the MitoFish database. The resulting ASV table was rarefied to an even depth for downstream analyses. Various quality control metrics, such as the number of reads, ASVs, and taxonomic assignments, were calculated to assess the reliability and reproducibility of the eDNA metabarcoding data.

### 2.3. Bioinformatic analysis and taxonomic assignment

The raw sequencing data were processed using the DADA2 pipeline in Rv4.0.3 (Callahan et al., 2016). Reads were filtered, dereplicated, denoised, and merged to infer amplicon sequence variants (ASVs) (Nearing et al., 2018). Chimeric ASVs were removed, and the remaining ASVs were assigned taxonomically using BLAST+ (Camacho et al., 2009) against the MitoFish database (Sato et al., 2018). BLAST hits were filtered and assigned to the lowest common ancestor using the MEGAN algorithm (Huson et al., 2016). ASVs not assigned to genus or below were removed. The ASV table was converted to a species-level occurrence matrix, rarefied to standardize sampling effort, and used to calculate alpha and beta diversity metrics with the vegan package (Dixon, 2003).

To evaluate the accuracy of eDNA metabarcoding, we compared the detected species with historical surveys and recent literature on the Ganjiang River (Guo et al., 2018; Hu et al., 2019). We calculated the proportion of true positives and false negatives, and examined potential factors contributing to false negatives, such as species rarity, body size, and habitat preference (Zhang et al., 2018).

### 2.4. Data analysis

We analyzed the eDNA metabarcoding data to assess the spatial and temporal patterns of fish diversity in the Ganjiang River. α-diversity indices (Chao1 richness estimator, Shannon diversity index, and Simpson diversity index) were calculated for each sample using the vegan package in Rv4.0.3 (Dixon, 2003), and compared between upstream and downstream sites, and between wet and dry seasons using Wilcoxon rank-sum tests. Spearman’s rank correlation was used to test the relationship between α-diversity and environmental variables. β-diversity was analyzed using NMDS based on Bray-Curtis dissimilarity, and the significance of spatial and temporal differences in fish community composition was tested using PERMANOVA and ANOSIM. RDA were used to identify the environmental factors influencing fish community structure. Indicator species analysis (Villeger et al., 2008) was performed to identify fish species significantly associated with different river sections or seasons, and Pearson’s correlation coefficients were calculated between the relative abundance of dominant species and environmental variables. All statistical analyses were conducted in Rv4.0.3 using the vegan, ggplot2, and indicspecies packages.

## 3. Results

### 3.1. eDNA sequencing output and fish species composition

A total of 114 water samples from the main stem of the Ganjiang River were collected and processed, yielding 12,345,678 valid sequences after quality filtering, with an average of 108,295 sequences per sample (range: 62,471-193,059). The concentrations of extracted eDNA samples ranged from 82.20 to 436.93 ng/μL in the wet season and from 21.67 to 411.37 ng/μL in the dry season, with overall higher concentrations in the wet season (Table S1). PCR amplification and gel electrophoresis results showed clear target bands for all samples, confirming the successful amplification of fish eDNA (Fig. S1).

The eDNA metabarcoding analysis revealed a diverse fish community in the Ganjiang River, with a total of 45 species detected, belonging to 7 orders, 14 families, and 38 genera (Table S2). The number of fish species detected per sample ranged from 12 to 37, with an average of 24 species. Temporally, the species richness was slightly higher in the wet season (41 species) compared to the dry season (39 species), with 36 species shared between the two seasons, indicating a relatively stable core fish community with some seasonal variations (Table S2).

### 3.2. Fish community composition and dominant species

The relative abundance of fish species varied spatially and temporally (Fig.2 A, B). In the wet season, the most abundant species were *Ctenopharyngodon idella* (28.3%), *Cyprinus carpio* (21.5%), *Hypophthalmichthys nobilis* (18.6%), and *Carassius auratus* (10.2%), which together accounted for 78.6% of the total reads. In the dry season, the dominant species shifted to *Cyprinus carpio* (32.1%), *Hypophthalmichthys nobilis* (22.4%), *Ctenopharyngodon idella* (15.3%), and *Silurus asotus* (8.6%), which contributed 78.4% of the total abundance. Spatially, the relative abundance of dominant species also varied among sites, with *Cyprinus carpio* being more abundant in the upstream sites, while *Ctenopharyngodon idella* and *Hypophthalmichthys nobilis* were more abundant in the downstream sites (Fig. 2C, D).

**Figure 2.**
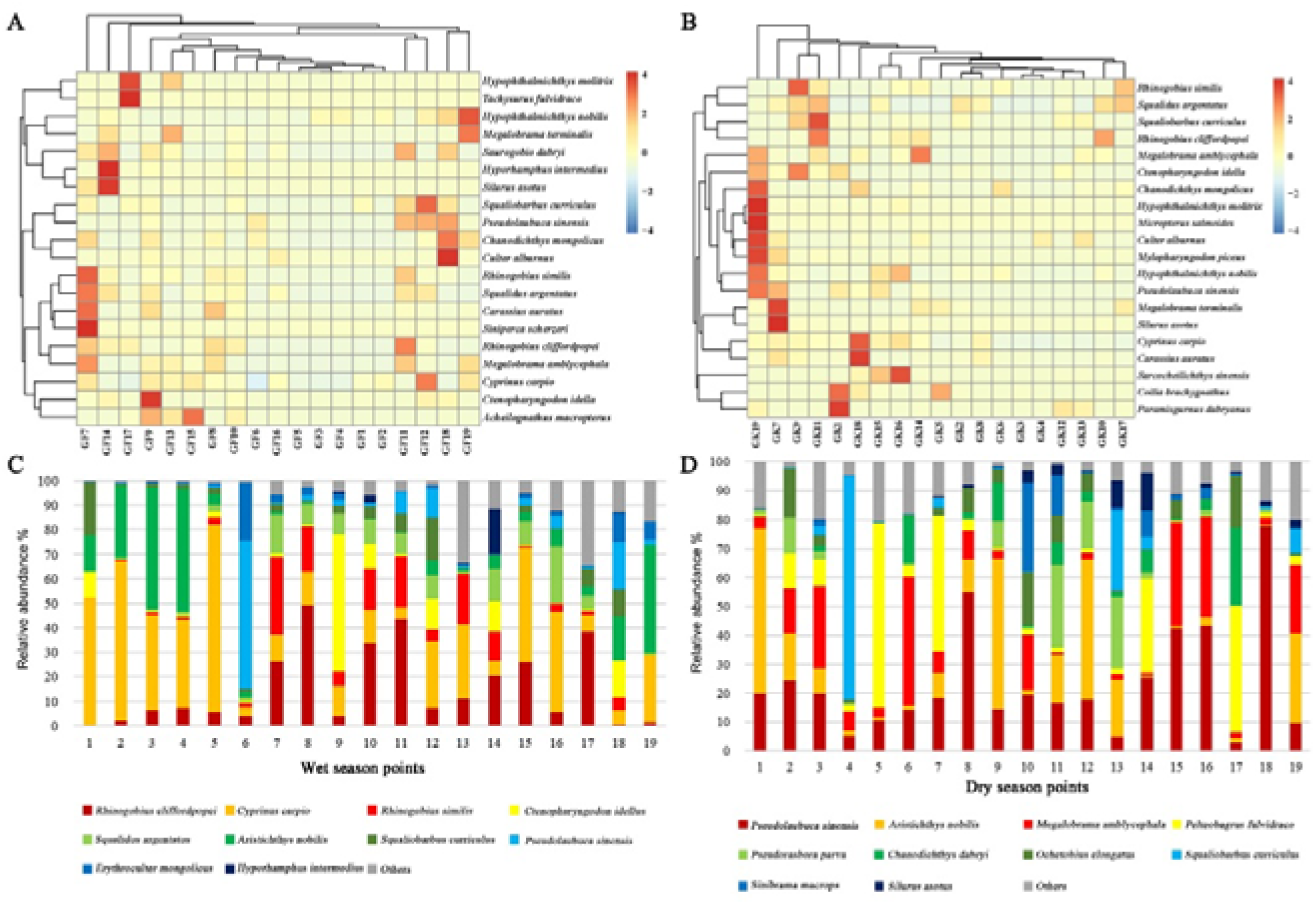
Relative abundance and distribution of dominant fish species in the Ganjiang River. (A) Heatmap showing the relative abundance at different sampling sites in the wet season; (B) Heatmap in the dry season; (C) The top 10 species in the wet season; (D) The top 10 species in the dry season.

Ten species were identified as dominant based on their relative abundance (>1%) and occurrence frequency (>50%) across all samples (Table 1). These species accounted for 92.1% of the total reads, with *Cyprinus carpio* being the most abundant (20.9%), followed by *Ctenopharyngodon idella* (15.0%), *Rhinogobius cliffordpopei* (12.3%), *Hypophthalmichthys nobilis* (9.5%), *Rhinogobius giurinus* (8.7%), *Squalidus argentatus* (6.5%), *Megalobrama amblycephala* (6.1%), *Squaliobarbus curriculus* (5.2%), *Pseudolaubuca sinensis* (3.6%), and *Chanodichthys mongolicus* (2.3%). The remaining 35 species were relatively rare, with relative abundances ranging from 0.01% to 0.9%.

**Table 1.**
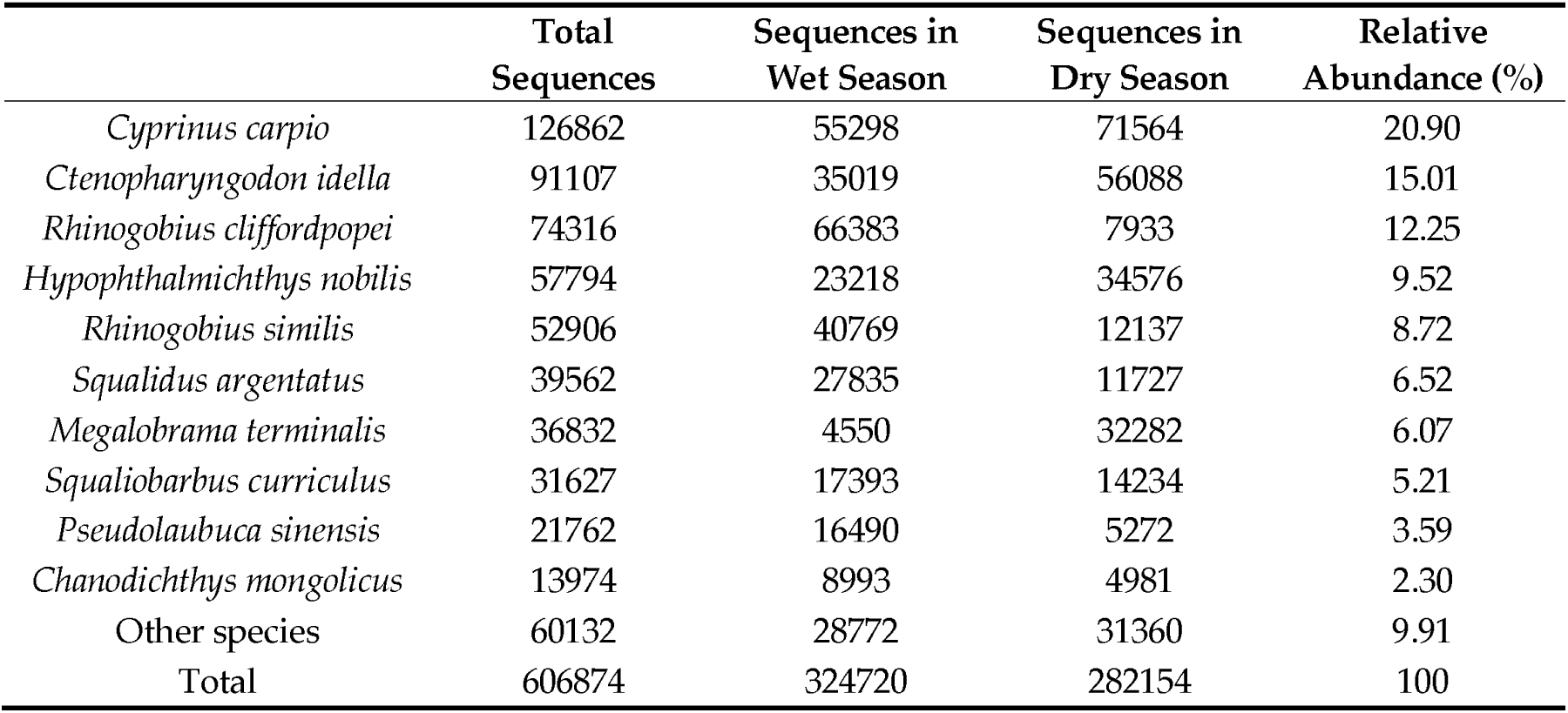
Relative abundance ratio of total sequence number in the two periods of the dominant species.

|  | Total<br>Sequences | Sequences in<br>Wet Season | Sequences in<br>Dry Season | Relative<br>Abundance (%) |
| --- | --- | --- | --- | --- |
| *Cyprinus carpio* | 126862 | 55298 | 71564 | 20.90 |
| *Ctenopharyngodon idella* | 91107 | 35019 | 56088 | 15.01 |
| *Rhinogobius cliffordpopei* | 74316 | 66383 | 7933 | 12.25 |
| *Hypophthalmichthys nobilis* | 57794 | 23218 | 34576 | 9.52 |
| *Rhinogobius similis* | 52906 | 40769 | 12137 | 8.72 |
| *Squalidus argentatus* | 39562 | 27835 | 11727 | 6.52 |
| *Megalobrama terminalis* | 36832 | 4550 | 32282 | 6.07 |
| *Squaliobarbus curriculus* | 31627 | 17393 | 14234 | 5.21 |
| *Pseudolaubuca sinensis* | 21762 | 16490 | 5272 | 3.59 |
| *Chanodichthys mongolicus* | 13974 | 8993 | 4981 | 2.30 |
| Other species | 60132 | 28772 | 31360 | 9.91 |
| Total | 606874 | 324720 | 282154 | 100 |

### 3.3. Spatial and temporal patterns of fish diversity

#### 3.3.1. Seasonal variations in fish diversity

The alpha diversity of fish communities in the Ganjiang River varied significantly between seasons (Fig. 3A). The average species richness per sample was significantly higher in the wet season (26.2 ± 4.5) than in the dry season (21.8 ± 3.8) (Wilcoxon rank-sum test, P < 0.001). The Shannon diversity index showed a similar trend, with significantly higher values in the wet season (2.51 ± 0.32) than in the dry season (2.23 ± 0.28) (P < 0.01).

**Figure 3.**
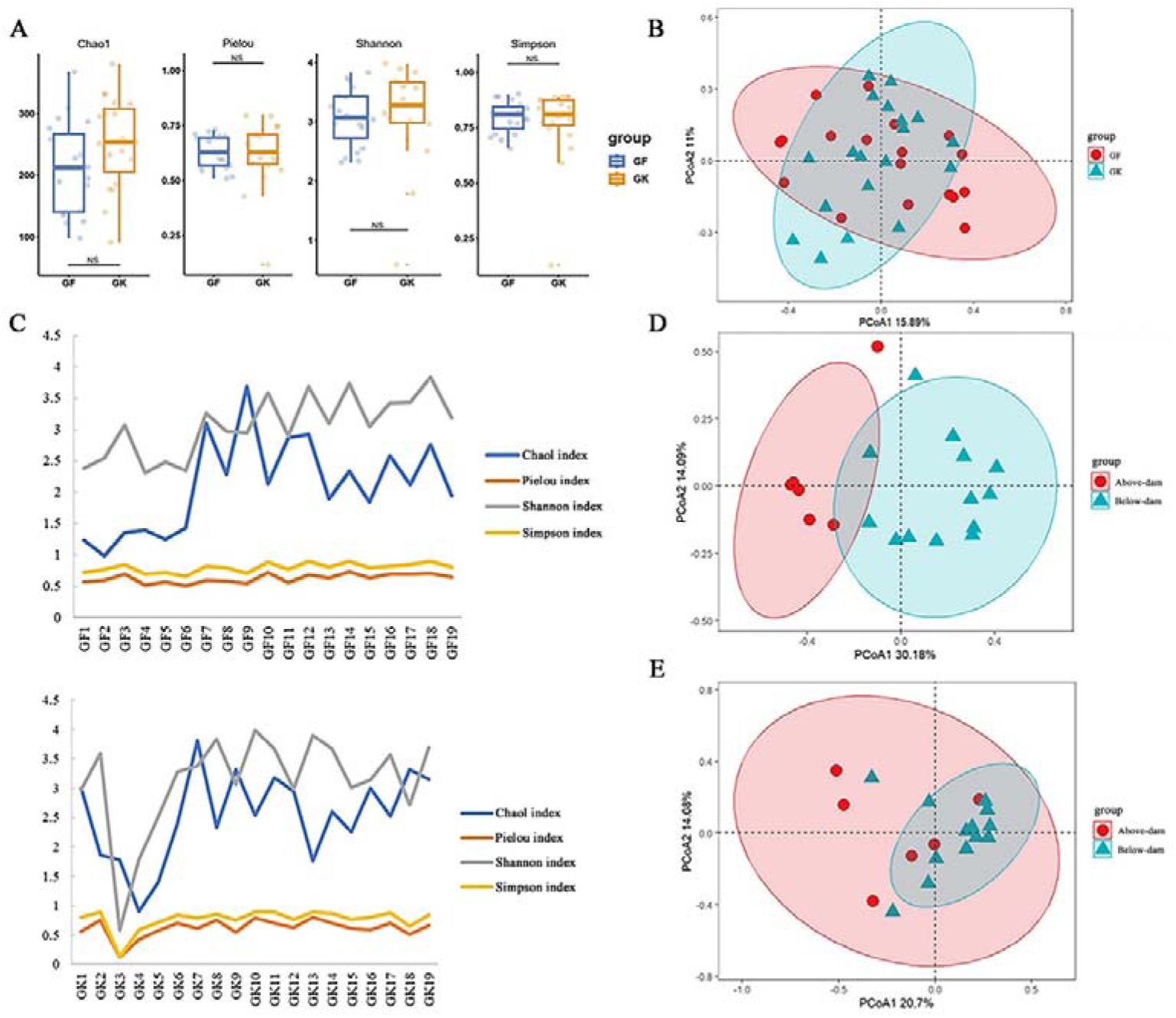
Spatio-temporal patterns of fish community diversity in the Ganjiang River based on eDNA. (A) Box plot of α diversity index and t test between groups in two periods of wet and dry; (B) PCoA map of fish assemblage composition in two periods of wet and dry; (C) Variation of α diversity index with point position in two periods of wetand dry; (D) PCoA map of fish assemblage composition before and after a dam in wet season; (E) PCoA map of fish assemblage composition before and after a dam in dry season.

The beta diversity analysis based on Bray-Curtis dissimilarity revealed significant differences in fish community composition between seasons (Fig. 3B). The PERMANOVA test showed that season had a significant effect on the community structure (R² = 0.052, P = 0.001), although it explained less variation than site (R² = 0.135, P = 0.001) (Table S3). The PCoA ordination plot clearly separated the wet and dry season samples along the first axis (PC1), which explained 21.5% of the total variation. The ANOSIM test further confirmed the significant differences in community composition between seasons (R = 0.376, P = 0.001).

#### 3.3.2. Spatial variations in fish diversity

The alpha diversity indices also exhibited significant spatial variations among sampling sites (Fig. 3C). In the wet season, the species richness and Shannon diversity index were highest at site 9 (downstream of the Wan’an Dam) and lowest at site 2 (within the Wan’an Reservoir). In the dry season, the highest values were observed at site 7 (downstream) for species richness and site 10 (downstream) for Shannon index, while the lowest values were found at site 4 (upstream) and site 3 (upstream), respectively. Overall, the fish diversity was higher in the downstream sites than in the upstream sites, especially those within the Wan’an Reservoir.

The beta diversity analysis further revealed significant differences in fish community composition between the upstream and downstream sites, particularly in the wet season (Fig. 3D, E). The PCoA ordination plot showed a clear separation of the upstream (sites 1-6) and downstream (sites 7-19) samples along the second axis (PC2), which explained 14.7% of the variation. The PERMANOVA test confirmed the significant effect of dam location on the community structure in both the wet season (R² = 0.266, P = 0.001) and the dry season (R² = 0.099, P = 0.045) (Table S4,5). The ANOSIM test also indicated significant differences in community composition between the upstream and downstream sites in the wet season (R = 0.512, P = 0.001) and the dry season (R = 0.276, P = 0.002).

### 3.4. Environmental factors influencing the spatiotemporal patterns of fish community structure

The RDA analysis also revealed significant spatial patterns in the relationships between environmental factors and fish communities. In the wet season, DO, TP, and PO43--P, had significant influences on the fish community structure (Fig. 4A). The upstream sites (1-6) were more strongly associated with DO, while the downstream sites (7-19) showed more variable relationships with different environmental factors. Most environmental factors were negatively correlated with the first RDA axis, except for PO43--P, and all factors except DO were negatively correlated with the second RDA axis.

**Figure 4.**
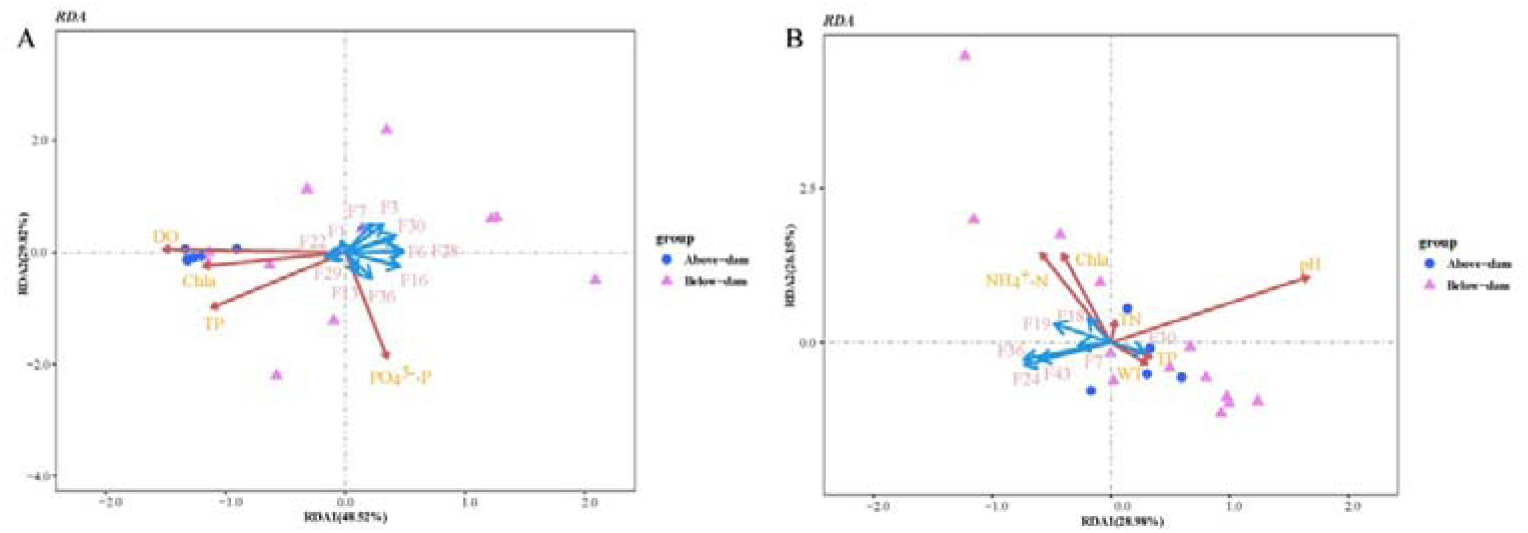
Redundancy analysis (RDA) revealing the relationships between fish communities and environmental variables in the Ganjiang River. (A) RDA analysis of environmental factors and fish communities in wet season; (B) RDA analysis of environmental factors and fish communities in dry season.

In the dry season, pH, NH4+-N, and chlorophyll a (Chla) were the most significant environmental factors influencing the fish community structure (Fig.4B). The upstream sites were more strongly associated with WT and TP, while the downstream sites were more influenced by pH, WT, and TP. pH, WT, TP, and TN were positively correlated with the first RDA axis, while TN, NH4+-N, and Chla were positively correlated with the second RDA axis.

## 4. Discussion

### 4.1. Fish biodiversity in the Ganjiang River revealed by eDNA

Our study provides the first comprehensive assessment of fish diversity in the Ganjiang River using eDNA metabarcoding, revealing a total of 45 species belonging to 7 orders, 14 families, and 38 genera. This species richness is lower than the historical records of over 80 fish species in the river basin (Hu et al., 2019; Huang et al., 2011).The discrepancy between eDNA and historical data may be due to several reasons. First, some rare or cryptic species might have been missed by eDNA sampling due to their low abundance or patchy distribution (Jerde et al., 2011). Second, the eDNA detection relies on the completeness and accuracy of reference databases, which may be limited for certain taxa or regions (Weigand et al., 2019). Third, the historical records might have included species from tributaries or lakes that were not covered by our sampling sites along the main stem. Nevertheless, the eDNA approach has demonstrated its effectiveness in detecting a wide range of fish taxa, including some important economic and ecological species, such as the four major Chinese carps (black carp, grass carp, silver carp, and bighead carp), and the endemic species like *Sinibrama macrops* and *Squaliobarbus curriculus* (Hu et al., 2022).

The fish community in the Ganjiang River was dominated by Cypriniformes, which accounted for 80% of the total species richness and over 90% of the total read abundance. This pattern is consistent with previous studies on the fish fauna of the Yangtze River basin, which is known for its high diversity and endemism of cyprinids (Yang et al., 2023). At the family level, Cyprinidae, Gobionidae, and Cobitidae were the most speciose and abundant groups, while other families like Bagridae, Siluridae, and Serranidae had lower diversity and abundance. The dominance of small-to-medium sized cyprinids and cobitids in the Ganjiang River may reflect the combined effects of habitat conditions, fishery activities, and sampling methods. The river contains many shallow and fast-flowing sections with gravel or sandy substrates, which favor the rheophilic and benthic species like gobionids and loaches (Shi et al., 2020). The intensive fishing pressure and habitat degradation may have depleted the populations of large-sized species such as common carp and culters, while the small-sized species are more likely to escape from the nets or traps (Zhang et al., 2019). The eDNA method is also known to have a higher detection probability for small or abundant species compared to large or rare species, due to the positive correlation between eDNA concentration and fish biomass or density (Doi et al., 2016).

Despite the relatively high species richness, the fish community in the Ganjiang River showed a low evenness and high dominance, with the top ten species accounting for over 90% of the total reads. The most abundant species were common carp, grass carp, bighead carp, and Wuchang bream, which are all commercially important and widely distributed in the Yangtze River basin (Yue et al., 2024). These species are known to be highly adaptable to various habitats and have high fecundity and growth rates, allowing them to maintain large populations in the altered river systems (Li et al., 2016a). In contrast, many native and endemic species, such as the Chinese sturgeon (*Acipenser sinensis*), Dabry’s sturgeon (*Acipenser dabryanus*), and Yangtze sturgeon (*Acipenser schrencki*), have become extremely rare or even extirpated from the Ganjiang River due to overfishing, habitat loss, and dam construction (Yang et al., 2023; Zhang et al., 2020). The eDNA metabarcoding could be a promising tool for monitoring the presence and distribution of these threatened species, as it has been successfully applied to detect the endangered Yangtze finless porpoise (*Neophocaena asiaeorientalis asiaeorientalis*) in the same river basin (Stewart et al., 2017).

In summary, our eDNA survey revealed a relatively high diversity but low evenness of fish species in the Ganjiang River, with the dominance of cyprinids and the rarity of some endemic and migratory species. These findings highlight the need for more effective conservation and management strategies to protect the fish biodiversity and ecosystem integrity of this important tributary of the Yangtze River. The eDNA approach provides a rapid and non-invasive tool for assessing the fish community composition and monitoring the population dynamics of key species, which can inform the conservation planning and evaluation of restoration efforts in the future.

### 4.2. Impact of cascade dams on fish community structure and diversity

Our study revealed significant spatial and temporal variations in the fish diversity and community composition of the Ganjiang River, which could be largely attributed to the impact of cascade dams. The α-diversity indices (species richness and Shannon index) and β-diversity analysis (NMDS and PERMANOVA) consistently showed that the fish community structure differed significantly between the upstream and downstream sites, especially in the wet season. The upstream sites within the impoundments of cascade dams had lower species richness and diversity, and were dominated by lentic or eurytopic species such as crucian carp, common carp, and silver carp. In contrast, the downstream sites had higher diversity and contained more rheophilic and migratory species such as Rhinogobius cliffordpopei, Rhinogobius giurinus, and Squalidus argentatus. These findings suggest that the cascade dams have fragmented the river system and created distinct habitats with different fish assemblages, which is consistent with previous studies on the ecological impacts of dams (Cheng et al., 2015; Liu et al., 2019a).

The dams can affect the fish community structure and diversity through multiple mechanisms. First, the dams block the migration routes of fish and isolate the populations, leading to the decline or loss of migratory species (Gao et al., 2019). Many fish species in the Ganjiang River, such as the four major Chinese carps, require long-distance migration between the river and lake habitats for spawning and nursery (Zhang et al., 2020). The construction of cascade dams has disrupted their life cycles and reduced their recruitment, as evidenced by the low abundance of these species in the impounded sections. Second, the dams alter the hydrological regimes and habitat conditions of the river, creating lentic environments with deeper water, slower flow, and finer substrates (Liu et al., 2024). These changes favor the proliferation of limnophilic species and the elimination of rheophilic species, resulting in the homogenization of fish communities (Ding et al., 2017). Third, the dams trap the sediments and nutrients in the reservoirs, reducing the downstream transport and degrading the water quality and habitat suitability for fish (Xu et al., 2013). The decreased turbidity and increased water clarity may also increase the predation pressure on fish eggs and larvae, further impacting the fish populations (Liu et al., 2019a).

The impact of cascade dams on fish diversity also varied seasonally, with more pronounced differences between upstream and downstream sites in the wet season than in the dry season. This could be related to the migration and spawning behaviors of fish in response to the hydrological cues. Many fish species in the Ganjiang River, such as the four major Chinese carps, rely on the rising water levels and increasing flows in the wet season to stimulate their upstream migration and spawning activities (Gao et al., 2016). However, the dams regulate the water discharge and weaken the natural flow pulses, which may disrupt the reproductive cycles of fish and reduce their diversity in the impounded sections (Yi et al., 2010). In the dry season, the water levels and flows are generally low and stable, and the fish are less active in migration or spawning. Therefore, the impact of dams on fish diversity may be less evident in this period.

Despite the overall negative impact of cascade dams on fish diversity, our study also found some potential refugia or hotspots for fish conservation in the Ganjiang River. For example, the sites immediately downstream of the dams (e.g., sites 7 and 9) had relatively high species richness and contained some unique species that were absent from other sites. These sites might benefit from the dam-induced flow regulation and nutrient release, which could enhance the habitat heterogeneity and productivity for fish (LI et al., 2016b). The tributaries and floodplain lakes connected to the main stem (e.g., sites 13 and 19) also harbored diverse fish assemblages and could serve as important spawning and nursery grounds for migratory species (Liu et al., 2019b). Therefore, protecting and restoring these critical habitats should be a priority for the conservation and management of fish diversity in the Ganjiang River.

In conclusion, our study demonstrated the significant impact of cascade dams on the fish community structure and diversity in the Ganjiang River, and highlighted the importance of considering the spatial and temporal variations in assessing the ecological consequences of dam construction. The eDNA metabarcoding approach provided a powerful tool for capturing the complex responses of fish communities to the environmental gradients and anthropogenic disturbances in this large river system. Our findings can inform the future planning and operation of cascade dams to minimize their negative impact on fish diversity and promote the sustainable management of fishery resources in the Yangtze River basin.

### 4.3. Spatio-temporal variations of fish community and underlying environmental drivers

Our study revealed distinct spatial and temporal patterns of fish community structure in the Ganjiang River, which could be attributed to the complex interplay of environmental factors and anthropogenic disturbances. The RDA analysis showed that water temperature, pH, dissolved oxygen, and nutrients (e.g., total phosphorus and ammonia nitrogen) were the key environmental variables shaping the fish assemblages, but their relative importance varied seasonally and spatially. In the wet season, dissolved oxygen and nutrients were the main drivers of fish community structure, with the upstream sites characterized by higher oxygen levels and lower nutrient concentrations compared to the downstream sites. This pattern may reflect the combined effects of natural longitudinal gradients and dam-induced alterations on the physicochemical conditions of the river (Vannote et al., 1980). The upstream reaches tend to have faster flow, shallower depth, and coarser substrates, which facilitate the aeration and oxygenation of water (Wetzel, 1975). The dams can further enhance the dissolved oxygen levels in the impounded sections by increasing the water residence time and promoting the phytoplankton production (Baumgartner et al., 2021). However, the dams can also trap the nutrients in the sediments and alter their downstream transport, leading to the oligotrophication of the river (Maavara et al., 2020).

In the dry season, water temperature, pH, and conductivity emerged as the primary environmental factors affecting the fish community structure, with the upstream sites showing higher pH and lower temperature and conductivity than the downstream sites. This pattern suggests that the fish assemblages are sensitive to the changes in water quality and thermal regimes during the low-flow period (Olden and Naiman, 2009). The upstream reaches are more influenced by the groundwater inputs and tributary inflows, which tend to have lower temperature, higher pH, and lower ionic concentrations (Poole and Berman, 2001). The dams can also alter the thermal stratification and water chemistry of the impounded sections, creating a lentic environment with higher temperature and conductivity (Preece and Jones, 2002). These environmental gradients can filter the fish species based on their physiological tolerances and habitat preferences, leading to the differentiation of fish communities along the river continuum (Kirk et al., 2021).

The temporal variations of fish community structure in the Ganjiang River could be related to the seasonal changes in hydrology, water quality, and biotic interactions. The PERMANOVA results indicated that the fish assemblages differed significantly between the wet and dry seasons, with higher dissimilarity and turnover in the wet season. This pattern may be driven by the increased river connectivity and habitat heterogeneity during the high-flow period, which facilitates the dispersal and recruitment of fish species (Shao et al., 2019). The rising water levels and faster currents in the wet season can also stimulate the spawning and migration of some rheophilic and potamodromous species, such as the Rhinogobius and Squalidus (Yu and Lee, 2002). In contrast, the dry season is characterized by lower water levels, slower flows, and higher stability, which may limit the movement and reproduction of fish (Li et al., 2023). The reduced connectivity and increased isolation of river segments can also intensify the biotic interactions, such as competition and predation, leading to the local exclusion or extirpation of certain species (Shao et al., 2019).

The environmental drivers of fish community structure in the Ganjiang River are likely to be influenced by multiple stressors, including dam construction, land use change, and climate variability (Dudgeon, 2019). The cascade dams can modify the natural flow regimes, sediment transport, and nutrient cycling, creating novel ecological conditions that favor some species while disadvantaging others (Chen et al., 2024). The land use activities, such as agriculture, urbanization, and deforestation, can increase the nutrient and sediment inputs into the river, leading to the eutrophication and siltation of aquatic habitats (Shan et al., 2023). The climate change, particularly the increased temperature and altered precipitation patterns, can affect the water availability, thermal stability, and biotic interactions in the river ecosystems (Ficke et al., 2007). These stressors can act synergistically or antagonistically on the fish communities, depending on their intensity, duration, and spatial extent (Birk et al., 2020).

To better understand the spatio-temporal variations of fish community structure in the Ganjiang River, future studies should integrate multiple approaches, such as long-term monitoring, trait-based analysis, and mechanistic modeling (Heino et al., 2021). The eDNA metabarcoding can provide a rapid and cost-effective tool for assessing the fish diversity and distribution across space and time, but it should be complemented by traditional sampling methods to validate the results and estimate the abundance and biomass of species (Hanfling et al., 2016). The trait-based analysis can help to identify the functional responses of fish assemblages to environmental gradients and anthropogenic disturbances, and predict their resilience and vulnerability to future changes (Vitule et al., 2016). The mechanistic modeling can simulate the complex interactions among the hydrological, biogeochemical, and ecological processes in the river basin, and evaluate the cumulative impacts of multiple stressors on the fish communities (Domisch et al., 2015). By combining these approaches, we can gain a more comprehensive and predictive understanding of the spatio-temporal dynamics of fish biodiversity in the Ganjiang River, and inform the conservation and management strategies for maintaining the ecological integrity and ecosystem services of this important tributary of the Yangtze River Basin.

### 4.4. Implications for conservation and management

Our findings have important implications for the conservation and management of fish diversity in the Ganjiang River basin. The construction of cascade dams significantly affected the spatial heterogeneity of fish communities in the Ganjiang River, resulting in significantly lower species diversity in the impounded sections compared to the downstream reaches. Therefore, it is necessary to integrate ecological, power generation, and flood control objectives in the planning and operation management of cascade development, optimize operation schemes, maintain suitable ecological flows, and mitigate the impact of habitat fragmentation on fish (Cheng et al., 2015; XIE, 2017). Consideration should be given to releasing appropriate ecological flows during the wet season to simulate natural flood pulses and promote fish migration and spawning (Li et al., 2020).

Nutrient concentrations (total phosphorus, ammonia nitrogen, etc.) are important environmental factors affecting the spatial and temporal distribution of fish communities in the Ganjiang River, reflecting the stress of agricultural non-point source pollution on water quality in the basin. Best management practices such as fertilizer control, riparian buffers, and vegetative filter strips should be adopted to control agricultural runoff from entering the river at the source and pathway, and alleviate the stress of eutrophication on fish and their food webs (Chen et al., 2014; Xia et al., 2006).

The cascade dams on the main stem of the Ganjiang River have blocked the migration of fish species, preventing them from completing their life history. There is an urgent need to construct fishways and other aquatic ecological corridors to restore the connectivity between the river and lakes, and provide migration pathways for economically important species such as the four major Chinese carps and rare species like the Chinese giant salamander (Shi et al., 2017).

Climate change and land use change will further exacerbate river habitat degradation and environmental stress, requiring integrated management from a watershed perspective. Strengthening the constraints of regional planning environmental impact assessment and watershed ecological protection red lines, and controlling development activities that are not conducive to fish diversity conservation, such as land reclamation from lakes and riverbank hardening, are crucial. Carrying out ecological restoration projects such as returning farmland to wetlands, wetland restoration, and river-lake connectivity, as well as improving soil and water conservation capacity and reducing sediment and non-point source pollution in the river, are also important measures (Albert et al., 2021; Fan et al., 2015).

Environmental DNA metabarcoding technology is expected to become a regular monitoring and early warning tool for fish diversity, providing timely and accurate baseline data and change information for watershed management agencies (Li et al., 2009; Wang et al., 2018). The monitoring scope should be further expanded to tributaries and lakes, and an integrated monitoring network for hydrology, water quality, and aquatic ecology should be developed and incorporated into the watershed ecological security early warning and decision support system.

These implications and recommendations can provide references for the conservation of fish diversity and management of cascade development in the Ganjiang River and other tributaries of the Yangtze River Basin. Future studies should further integrate research on environmental DNA, functional traits, food webs, and genetic diversity to reveal the ecological adaptation mechanisms of fish communities under cascade disturbances, and combine scenario analysis and land-sea integrated models to predict the impacts of future environmental changes and develop more comprehensive and effective conservation strategies. At the same time, efforts should be made to translate scientific research results into management decisions, promote the transition of watershed integrated management from engineering-based approaches to ecosystem-based management, and achieve harmonious coexistence between humans and nature.

## 5. Conclusions

In this study, we used environmental DNA (eDNA) metabarcoding to assess the spatiotemporal patterns of fish diversity and community structure in the Ganjiang River, a major tributary of the Yangtze River, in relation to cascade hydropower development. Our results revealed significant spatial and temporal variations in fish diversity, with lower species richness and distinct community composition in the impounded sections compared to the downstream reaches, especially during the wet season. The fish assemblages were strongly influenced by environmental factors such as water temperature, pH, dissolved oxygen, and nutrients, which exhibited complex interactions with dam operation and land use practices. Our findings highlight the importance of integrating ecological considerations into the planning, construction, and operation of cascade hydropower projects to mitigate their adverse impacts on fish diversity. We recommend adopting ecologically friendly dam operation schemes, such as releasing environmental flows to maintain river connectivity and simulate natural flow pulses, controlling agricultural non-point source pollution, constructing fishways and aquatic ecological corridors, strengthening watershed-scale integrated management, and conducting regular monitoring and early warning using eDNA tools. This study demonstrates the effectiveness of eDNA metabarcoding in assessing the responses of fish communities to anthropogenic disturbances and environmental gradients in large river systems. By providing a comprehensive and high-resolution view of fish diversity patterns, eDNA-based biomonitoring can inform more targeted and effective conservation and management strategies. As the global demand for clean energy continues to grow, it is crucial to strike a balance between sustainable hydropower development and the conservation of freshwater biodiversity. The Ganjiang River, with its rich fish fauna and extensive cascade hydropower system, provides a valuable case study for understanding and managing the complex interactions between dams and fish communities in large river basins. Our study contributes to the growing body of knowledge on the ecological consequences of hydropower development and highlights the need for science-based, adaptive, and participatory approaches to achieving sustainable river basin management in the face of multiple stressors and competing demands.6.

## Supplementary Materials

Figure S1: PCR amplification and gel electrophoresis results showed clear target bands for all samples, confirming the successful amplification of fish eDNA; Table S1: Environmental DNA concentration at different sampling sites; Table S2: Fish information of Ganjiang River main stream was obtained based environmental DNA technique; Table S3: The parameters and two Permanova analysis calculated value; Table S4: The calculated values of each parameter were analyzed by Permanova in wet season; Table S5: The calculated values of each parameter were analyzed by Permanova in wet season.

## Institutional Review Board Statement

This study did not involve humans or animals and only used environmental water samples. Therefore, IRB approval was not required.

