## supplementary materials for "Assessing fish biodiversity and community structure in the Ganjiang River using eDNA metabarcoding"

**Assessing fish biodiversity and community structure in the Ganjiang River using eDNA metabarcoding: Impact of cascade hydropower development**

Total of 6 pages, not including the cover page, with 1 figures, 5 table.

Supplementary Materials includes the following:

**Figure S1.** PCR amplification and gel electrophoresis results showed clear target bands

for all samples, confirming the successful amplification of fish eDNA .

**Table S1**. Environmental DNA concentration at different sampling sites.

**Table S2**. Fish information of Ganjiang River main stream was obtained based environmental DNA technique.

**Table S3**. The parameters and two Permanova analysis calculated value.

**Table S4**. The calculated values of each parameter were analyzed by Permanova in wet season.

**Table S5**. The calculated values of each parameter were analyzed by Permanova in wet season.

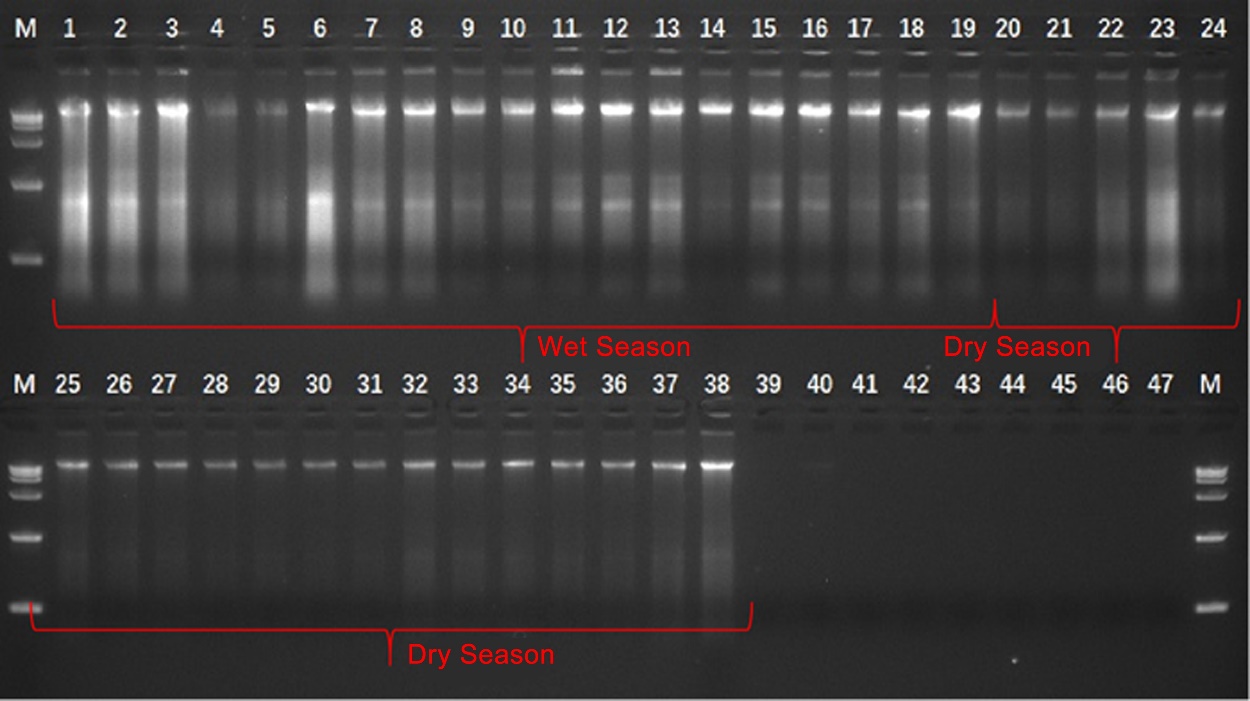

Figure S1. PCR amplification and gel electrophoresis results showed clear target bands for all samples, confirming the successful amplification of fish eDNA .

Tab. S1Environmental DNA concentration at different sampling sites

| Samples during the wet season | DNA concentration ng/uL | Samples during the dry season | DNA concentration ng/uL |
| --- | --- | --- | --- |
| 1 | 436.93 | 1 | 73.97 |
| 2 | 366.47 | 2 | 64.50 |
| 3 | 309.53 | 3 | 148.30 |
| 4 | 98.83 | 4 | 411.37 |
| 5 | 91.43 | 5 | 92.90 |
| 6 | 392.57 | 6 | 49.23 |
| 7 | 247.87 | 7 | 31.43 |
| 8 | 245.90 | 8 | 37.93 |
| 9 | 112.30 | 9 | 25.97 |
| 10 | 122.17 | 10 | 29.37 |
| 11 | 179.13 | 11 | 21.67 |
| 12 | 188.50 | 12 | 28.03 |
| 13 | 261.93 | 13 | 60.80 |
| 14 | 82.20 | 14 | 43.37 |
| 15 | 170.73 | 15 | 48.87 |
| 16 | 184.40 | 16 | 39.20 |
| 17 | 145.07 | 17 | 42.10 |
| 18 | 240.17 | 18 | 77.67 |
| 19 | 146.57 | 19 | 124.10 |

Tab. S2 Fish information of Ganjiang River main stream was obtained based on environmental DNA technique

| Order | Family | Genus | Species | Wet season | Dry season |
| --- | --- | --- | --- | --- | --- |
| Cypriniformes | Acheilognathidae | *Acheilognathus* | *Acheilognathus macropterus* | + | + |
|  |  | *Rhodeus* | *Rhodeus ocellatus* | + | + |
|  | Cyprinidae | *Carassius* | *Carassius auratus* | + | + |
|  |  | *Cyprinus* | *Cyprinus carpio* |  | + |
|  | Xenocyprididae | *Chanodichthys* | *Chanodichthys dabryi* | + | + |
|  |  |  | *Chanodichthys mongolicus* | + | + |
|  |  | *Ctenopharyngodon* | *Ctenopharyngodon idella* | + | + |
|  |  | *Culter* | *Culter alburnus* | + | + |
|  |  | *Distoechodon* | *Distoechodon tumirostris* |  | + |
|  |  | *Elopichthys* | *Elopichthys bambusa* |  | + |
|  |  | *emiculter* | *Hemiculter leucisculus* | + | + |
|  |  | *Hypophthalmichthys* | *Hypophthalmichthys molitrix* | + | + |
|  |  |  | *Hypophthalmichthys nobilis* | + | + |
|  |  | *Megalobrama* | *Megalobrama amblycephala* | + | + |
|  |  |  | *Megalobrama terminalis* | + | + |
|  |  | *Mylopharyngodon* | *Mylopharyngodon piceus* | + | + |
|  |  | *Ochetobius* | *Ochetobius elongatus* |  | + |
|  |  | *Parabramis* | *Parabramis pekinensis* | + | + |
|  |  | *Pseudobrama* | *Pseudobrama simoni* | + | + |
|  |  | *Pseudolaubuca* | *Pseudolaubuca engraulis* | + |  |
|  |  |  | *Pseudolaubuca sinensis* | + | + |
|  |  | *Sinibrama* | *Sinibrama macrops* | + | + |
|  |  | *Squaliobarbus* | *qualiobarbus curriculus* | + | + |
|  |  | *Xenocypris* | *Xenocypris davidi* |  | + |
|  |  | *Zacco* | *Zacco acanthogenys* |  | + |
|  | Gobionidae | *Sarcocheilichthys* | *Sarcocheilichthys parvus* | + |  |
|  |  |  | *Sarcocheilichthys sinensis* |  | + |
|  |  | *Saurogobio* | *Saurogobio dabryi* | + | + |
|  |  |  | *Saurogobio gracilicaudatus* | + |  |
|  |  | *Squalidus* | *Squalidus argentatus* | + | + |
|  |  | *Microphysogobio* | *Microphysogobio elongatus* | + | + |
|  |  | *Hemibarbus* | *Hemibarbus labeo* | + | + |
|  |  | *Platysmacheilus* | *Platysmacheilus longibarbatus* | + |  |
|  |  | *Pseudorasbora* | *Pseudorasbora parva* |  | + |
|  | Botiidae | *Parabotia* | *Parabotia fasciatus* | + |  |
|  | Cobitidae | *Paramisgurnus* | *Paramisgurnus dabryanus* | + | + |
| Siluriformes | Siluridae | *Silurus* | *Silurus asotus* | + | + |
|  | Bagridae | *Tachysurus* | *Tachysurus fulvidraco* | + | + |
| Centrarchiformes | Centrarchidae | *Micropterus* | *Micropterus salmoides* |  | + |
|  | Sinipercidae | *Siniperca* | *Siniperca scherzeri* | + |  |
| Gobiiformes | Gobies | *Rhinogobius* | *Rhinogobius cliffordpopei* | + | + |
|  |  |  | *Rhinogobius similis* | + | + |
| Clupeiformes | Engraulidae | *Coilia* | *Coilia brachygnathus* | + | + |
| Synbranchiformes | Synbranchidae | *Monopterus* | *Monopterus albus* |  | + |
| Beloniformes | Hemiramphidae | *Hyporhamphus* | *Hyporhamphus intermedius* | + | + |

Tab. S3 The parameters and two Permanova analysis calculated value

|  | Factor | df | SS | MS | F | R² |
| --- | --- | --- | --- | --- | --- | --- |
| Season | 1 | 0.5643 | 0.5643 | 2.0799 | 0.0546 | 0.0130 |
| Residuals | 36 | 9.7678 | 0.2713 |  | 0.9454 |  |
| Total | 37 | 10.3322 |  |  |  |  |

Tab. S4 The calculated values of each parameter were analyzed by Permanova in wet season

|  | Factor | df | SS | MS | F | R² |
| --- | --- | --- | --- | --- | --- | --- |
| Season | 1 | 1.2105 | 1.2105 | 6.1601 | 0.2660 | 0.0010 |
| Residuals | 17 | 3.3407 | 0.1965 |  | 0.7340 | 17 |
| Total | 18 | 4.5512 |  |  |  |  |

Tab. S5 The calculated values of each parameter were analyzed by Permanova in wet season

|  | Factor | df | SS | MS | F | R² |
| --- | --- | --- | --- | --- | --- | --- |
| Season | 1 | 0.5169 | 0.5169 | 1.8698 | 0.0991 | 0.0450 |
| Residuals | 17 | 4.6997 | 0.2765 |  | 0.9009 | 17 |
| Total | 18 | 5.2166 |  |  |  |  |
